# Development of Microbiome Biomarkers for IgA: a Joint Modeling Approach

**DOI:** 10.1101/2021.04.15.439964

**Authors:** Rudradev Sengupta, Olajumoke Evangelina Owokotomo, Ziv Shkedy

## Abstract

Our aim in this study is to develop predictive microbiome biomarkers for intestinal IgA levels. In this article, a operational taxonomic units(OTU)-specific (family-specific) and time-specific joint model is presented as a tool to model the association between OTU (or family) and biological response (measured by IgA level) taking into account the treatment group (Control or PAT) of the subjects. The model allows detecting OTUs (families) that are associated with the IgA; for some OTUs (families), the association is driven by the treatment while for others the association reflects the correlation between the OTUs (families) and IgA.The results of the analysis reveal that: (1) the observed diversity of S24-7 family can be used as a biomarker to classify samples according to treatment group for days 6 and 12; (2) the treatment effect induces the corrlelation between the S24-7 diversity and the IgA level at day 20; (3) The OTUs that are identified to be significantly differentially abundant (FDR level of 0.05) between the two treatment groups for days 12 and 20 are all part of the S24-7 family, although most of the differentially abundant ones at day 1 are from the Lactobacillaceae family; (4) only the Lachnospiraceae family diversity at day 6, and 20 can be used as predictive biomarker for the IgA level at day 20; (5) New.ReferenceOTU513, correlated with the IgA level at day 20, since day 12, belongs to the Lachnospiraceae family and all other OTUs among the top 10 significantly associated OTUs at day 20 are from the S24-7 family; (6) the observed alpha diversity at day 6 is significantly differentially abundant and can be used as predictive biomarker for IgA level at day 20.

## 1 Introduction

In the past few years, there has been an increase in interest to study the association between compositions of microbial communities and different diseases (Kostic et al., 2014; Parekh et al., 2015; John and Mullin, 2016; Young, 2017; Wang et al., 2017). Although methods to identify different compositions of microbial communities across diseases levels are well developed, the development of new methods to identify microbiome biomarkers, i.e., methods to model the association between the microbiome variables and a clinical endpoint, is of primary interest.

In the current study, we present a new method that can be used to identify high dimensional microbiome biomarkers for the immune system which is measured using intestinal Immunoglobulin A (IgA) levels, taking into account a possible treatment effect on both variables. We present a joint model (Perualila-Tan et al., 2016; Perualila et al., 2016) for the microbiome biomarker and IgA that allows to include the treatment (and possibly other confounders) in the model as a co-variate(s). As a case study, we use an experiment where germ-free mice were conventionalized with a normal or antibiotic-perturbed microbiota with the aim of understanding the effect of antibiotic administration on the intestinal microbiota and host immunity (Ruiz et al., 2017). The dataset consists of 15 murine subjects and 355 OTUs, with representation > 0.01% in relative abundance. Longitudinal microbiome measurements (OTU counts) and longitudinal Immunoglobulin A (IgA) data were available at 7 common timepoints for all the subjects randomized into the two treatment groups. groups. For the analysis presented in this paper we used the microbiome measuerments in the first 4 times points (day 1, 6, 12 and 20) and the IgA level in day 20. Our goal is to link between the microbiome measurments and the IgA, taking into account that the treatment may influence both microbiome and IgA data. The time-specific joint mode presented in this paper allows to model two types of relationships between IgA level and OTU relative abundance: (1) an association which is driven by the treatment effect and (2) an association reflecting the correlation between the OTUs and IgA. Although we have longitudinal measurements available, in this article, we only focus on identifying potential microbiome biomarkers that can serve as indicator of the IgA response at a given timepoint and not on modeling the mean evolution of microbiome variables and IgA levels over time. The proposed joint model is flexible in the sense that it can accommodate microbiome measuerments in different resolutions. For the analysis presented in this paper we used the family level richness i.e., the number of OTUs belonging to the family with non zero counts and the relative abundance as microbiome covariates which potentially can be used as biomarkers for IgA.

This paper is organised as follows: In Section 2, we introduce the data setting in the transPAT experiment. In Section 3, the joint modeling approach is formulated while in Section 4 the model is applied to both family level and otu level data. Finally, we discuss the results in Section 5.

## 2 Data

### 2.1 transPAT

In this study, 15 murine subjects were randomized into two treatment groups. The main motivation is to investigate if a single pulsed antibiotic treatment (PAT) course at a stage early in life, can cause withstanding alterations to the intestinal microbiota (Ruiz et al., 2017). Initially a set of germ-free mice are divided into two groups with one group receiving tylosin as treatment and the other receiving placebo. The microbiota from these subjects were transplanted in another new set of 15 germ-free mice with one group serving as the control group consisting of 8 mice while the other group consisting of 7 mice are recipients of pulsed antibiotic treatment (PAT) perturbed microbiota. Subjects were followed over time and both their fecal IgA levels as well as the sequence counts data for 355 Operating Taxonomic Units (OTUs) with representation > 0.01% in relative abundance were measured over different timepoints. Hence, for each timepoint, we have the following data set-up:
IgA level (*Y*):

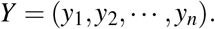

Microbiome data (**X**):

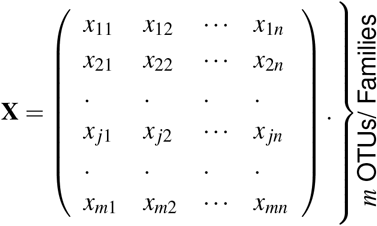

Binary indicator for treatment group (*Z*):

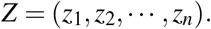

Figure 1a and Figure 1b show the longitudinal measuerments of IgA and the boxplot of IgA at day 20. Figure 1c shows an example OTU (264734) wih the change in relative abundance over time. For the analysis in this paper we use the relative abundance at each time point separately. As measure for microbiome activity at a family level we use the family level richness, shown in Figure 1d. The family level richness is the number of OTUs, belonging to a specific family, that have non zero counts. Figure 1e shows the richness profiles of the *Dehalobacteriaceae* family and reveal a non active family, in contrast with the *S24-7* family shown in Figure 1d. The issue of active and non active families is discussed further in Section 2 in the appendix. Figure 1f shows the *α* diversity (Morgan and Huttenhower, 2012) profiles for the study which is the richness at a Kingdom level.

**Figure 1:**
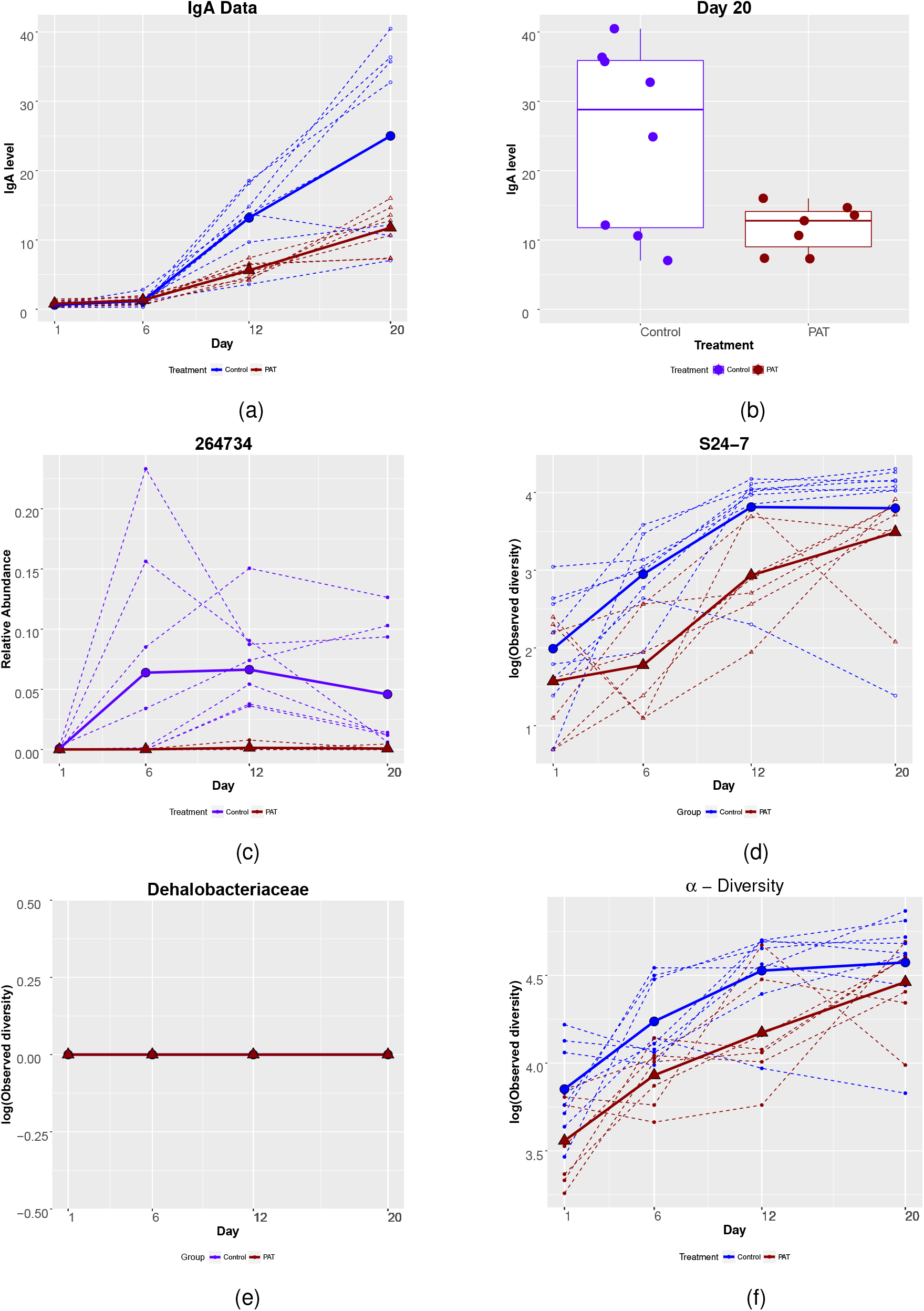
(a) The IgA level over time; (b) Boxplot of the IgA level of 15 mice by Treatment for Day 20; (c) Example of an OTU over time; (d) Active *S24-7* family over time; (e) Non-active *Dehalobacteriaceae* family over time; (f) Overall *α* diversity, in log scale, over time. For all the figures dashed lines and solid lines represent subject profiles and mean profiles, respectively.

## 3 Modeling Approach

### 3.1 A Joint Model for Microbiome Measurements and IgA

Our aim is to identify microbiome biomarkers for IgA. For this purpose we formulated a joint model for the microbiome variables and IgA. Let **X** be a *m × n* timepoint specific microbiome measurements matrix in which columns represent subjects and rows represent microbiome variables. The microbiome variables depend on the resolution at which the model is fitted. For OTU specific model, *X_ji_, i* = 1, *… n, j* = 1*,…m* is the measurement for the *j*th OTU of the *i*th subject. If the model is fitted at a family level, *X_ji_* represents the richness of the *j*th family. note that if the model is applied to alpha diversity, **X** is a vector for which the *i*th entry corresponds to the alpha diversity of the *i*th subject. Let *Y_i_* denote the IgA level for the *i^th^* sample. The treatment group indicator is denoted by *Z* that takes a value of one (*z_i_* = 1) or zero (*z_i_* = 0) if the *i^th^* subject belongs to PAT or control group, respectively. Schematically, the OTU-specific and timepoint-specific joint model is presented in Figure 2.

**Figure 2:**
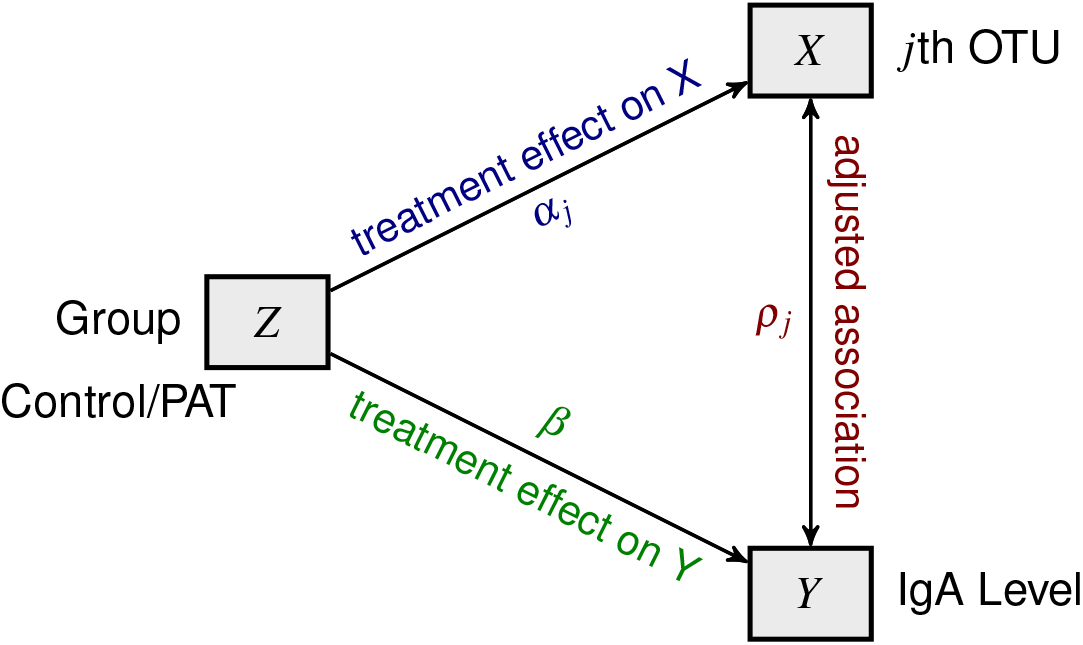
Illustration of the joint model for the microbiome measurements and IgA.

For a given timepoint, the OTU-specific joint model also allows us to test which OTU is differentially abundant and which OTU is predictive for the IgA measurement, taking into account a possible effect of the treatment on the two variables. Following Perualila *et al.* (2016a, 2016b) the joint model is formulated as follows:

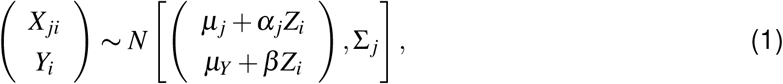

where the OTU-specific covariance matrix, Σ _*j*_, is given by,

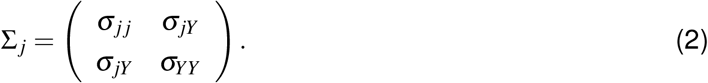

The parameters *α_j_* and *β* represent the treatment effects for the *j*th OTU and IgA data, respectively and *µ _j_* and *µ_Y_* are the average relative abundance for the *j*th OTU and the average of the IgA data, respectively, for mouse in the PAT group.

Thus, the OTU-specific association with the response can be obtained using adjusted association (Buyse and Molenberghs, 1998; Perualila et al., 2016; Perualila-Tan et al., 2016), a coefficient that is derived from the covariance matrix, Σ _*j*_,

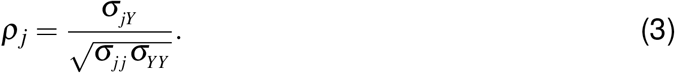

Indeed, *ρ_j_* = 1 indicates a deterministic relationship between the relative abundance of the *j*th OTU and the IgA response after accounting for the treatment effect.

Although, the joint model formulated in (1) is specified for the OTU level data, it can be fitted to any resolution of the data. For example, if **X** consists family level richness measurements the joint model can be used to identify families for which the microbiome activity can be used as biomarker to the immunological response. The same holds if the primary analysis is done based on alpha diversity.

### 3.2 Inference

As mentioned in section 3.1, the model allows testing for differentially abundant OTUs. Hence, for each microbiome variable, we test the hypothesis

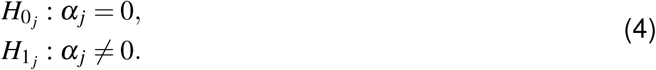

There are *m* null hypotheses are to be tested. Therefore, an adjustment for multiple testing should be applied. Throughout this paper, we apply the FDR approach proposed by Benjamini and Hochberg (1995). Furthermore, the joint model allows us to test whether the relative abundance of the OTUs and the IgA are correlated. Thus, in addition to the hypothesis in (4), one needs to test the hypothesis

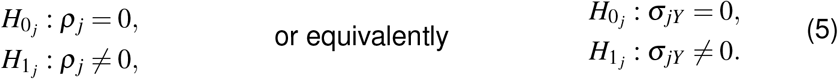

Note that under the null hypothesis, the covariance matrix of the joint model in (1) is reduced to

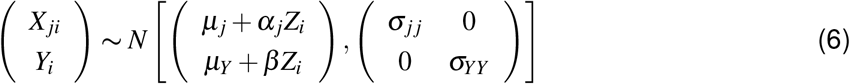

Consequently, the inference for the adjusted association can be done based on a likelihood ratio test by comparing the two models as specified by (1) and (6).

## 4 Results

The joint model specified in (1) was fitted to the transPAT data using different resolutions. In Section 4.1 we present the results for the analysis when family level richness is used as a biomarker while in Section 4.2 we present the results obtained for OTU level data. For the first analysis, log(richness) is used as a measure for microbiome activity at a family level and for the later, the relative abundance is used.

### 4.1 Analysis at Family Level

Observed *α*-diversity for a sample at a particular timepoint is the number of OTUs with non-zero counts at that particular timepoint and for that particular subject. We defined a new timepoint specific family-level index of *α*-diversity, as the number of active OTUs which belong to that family. For the *S24-7* family, the control group is dominated by the PAT group with respect to the observed diversity, across all the timepoints (Figure 1d). The joint model was fitted to the data for log(family-level richness) & IgA and significant families were identified based on treatment effect at 5% significance level. The families with more than 70% 0s per treatment group per timepoint were excluded and multiplicity adjustment was done accordingly (Benjamini and Hochberg, 1995).

#### 4.1.1 A Joint Model for the Richness of S24-7 Family and IgA

Table 1 displays the results for the *S24-7* family. Note that though this family was not significantly differentially abundant on day 1, it was found to be significant at day 6. However, it again became insignificant at later timepoints, day 12 and day 20. In Table 1, *α* estimates the difference in number of active OTUs, belonging to the *S24-7* family between the two treatment groups. Figure 3 and Table 1 show that initially this difference was small at day 1, then at day 6 1nd day 12 the control group had more number of active OTUs, belonging to this family and at day 20 the abundance in PAT group increased to make this difference small again.

**Table 1:** Parameter estimates from the model for the *S24-7* family at different timepoints.

| Day | S24-7 Family |  |  |  |  |  |
| --- | --- | --- | --- | --- | --- | --- |
| | $\alpha$ | $p(\alpha)$ | adj-p($\alpha$) | $\rho$ | $p(\rho)$ | adj-p($\rho$) |
| 1 | -0.42 | 0.30 | 0.49 | -0.41 | 0.10 | 0.49 |
| 6 | -1.17 | 0.00 | 0.00 | -0.03 | 0.92 | 0.99 |
| 12 | -0.88 | 0.01 | 0.07 | 0.40 | 0.11 | 0.22 |
| 20 | -0.31 | 0.48 | 0.77 | 0.53 | 0.03 | 0.05 |

**Figure 3:**
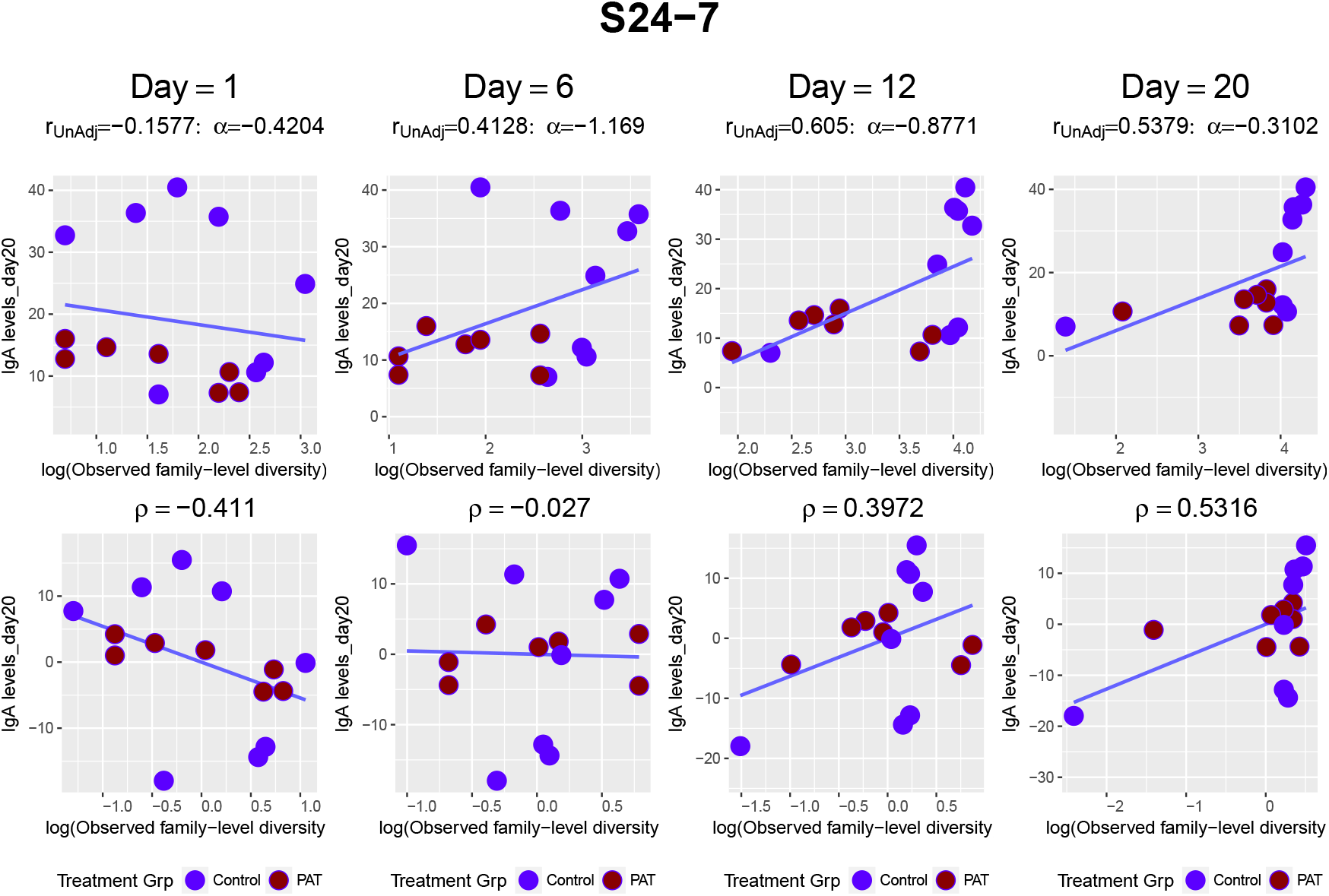
*S24-7* family over time.

Note that family level richness for the *S24-7* family, is found to be negatively associated with the IgA level at day 1 and day 6 which is in contrast with the positive association at later stages (day 12 and day 20). However, none of the associations are found to be significant.

#### 4.1.2 Other Families

Two families, namely *Erysipelotrichaceae* and *Verrucomicrobiaceae* >, are found to be significant at day 1 based on the treatment effect while at day 6 only the *S24-7* family is found to be significant. None were significant at day 12 and day 20.

When the adjusted association is explored, few new families are found to be important. No family was significant at day 1 and and day 12 whereas at both day 6 and day 20, there are at least three significant families. Among these families *Lachnospiraceae* (Figure S2 in supplementary appendix) is found to be significant at both day 6 and day 20 (Figure S3 in supplementary appendix). Table 2 displays the results for all the families when the observed family level richness and the IgA level were jointly modeled. Similar analysis was done for other measures (Chao1 and Shannon index) of richness (Chao, 1984; Shannon, 1948) and the results are available in the appendix.

**Table 2:**
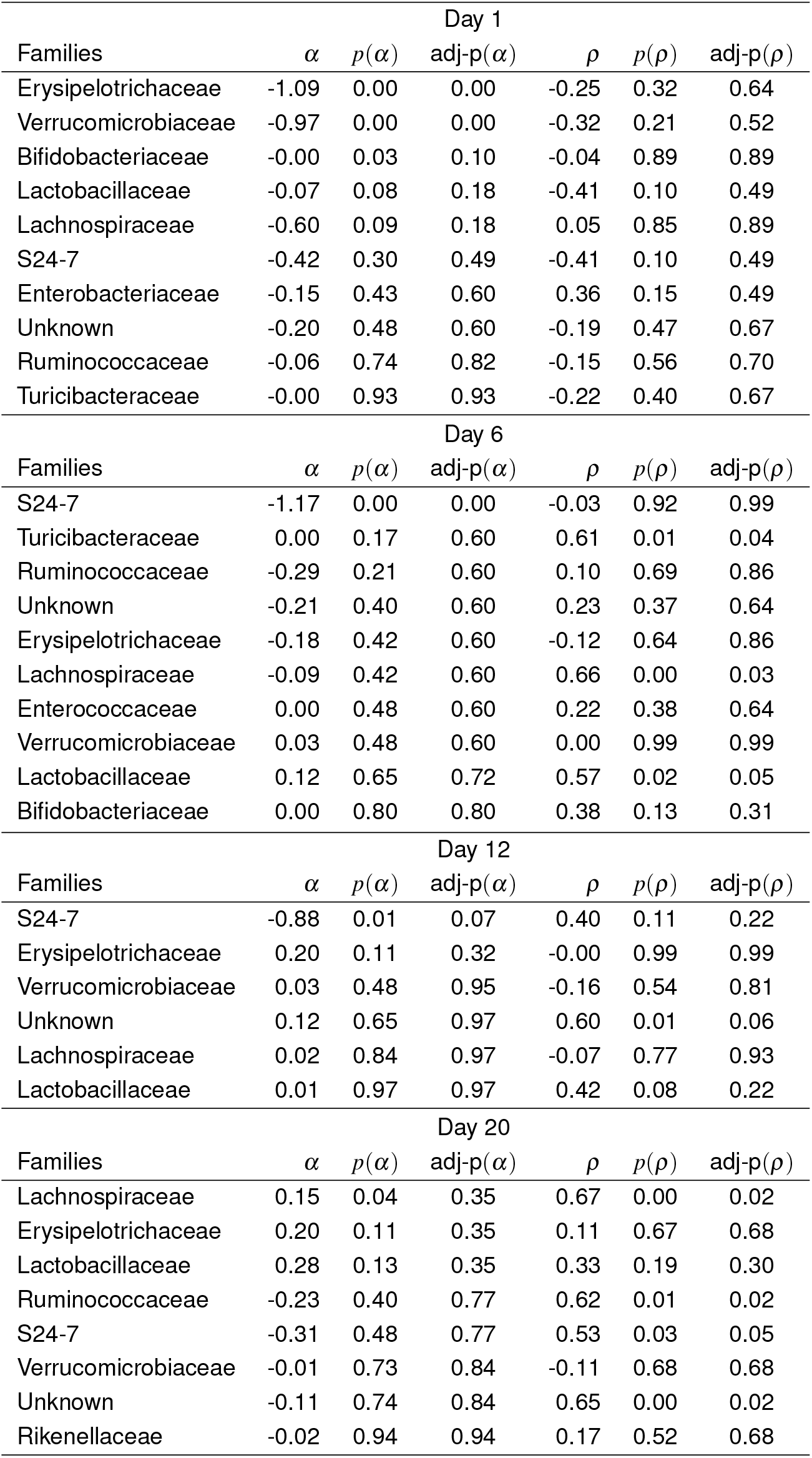
Parameter estimates from the model for all the families at different timepoints. Results are sorted according to the adjusted p-values for the treatment effect *α*.

### 4.2 Analysis at OTU Level

In Figure 4 (a,b,c,d), we plot the −log10 raw p-values of the adjusted association in x-axis and −log10 raw p-values of the treatment effect on microbiome in y-axis. The OTUs situated in the upper corners are differentially abundant between groups and those that are outlying with respect to the x-axis are OTUs that are correlated with the IgA level. Hence, we are interested on those that are lying in the upper right corner which are OTUs that are differentially abundant and have high adjusted association with IgA and could potentially serve as a predictive biomarker for the IgA level while OTUs in the upper left corner (e.g. 262095 at day 1, New.ReferenceOTU220 at day 12 - Figure 5) are differentially abundant but are conditionally independent of the IgA level. The OTUs at the bottom right (e.g. 221429 at day 12, 276629 at day 20 - Figure 6) have high adjusted association but are not differentially abundant.

**Figure 4:**
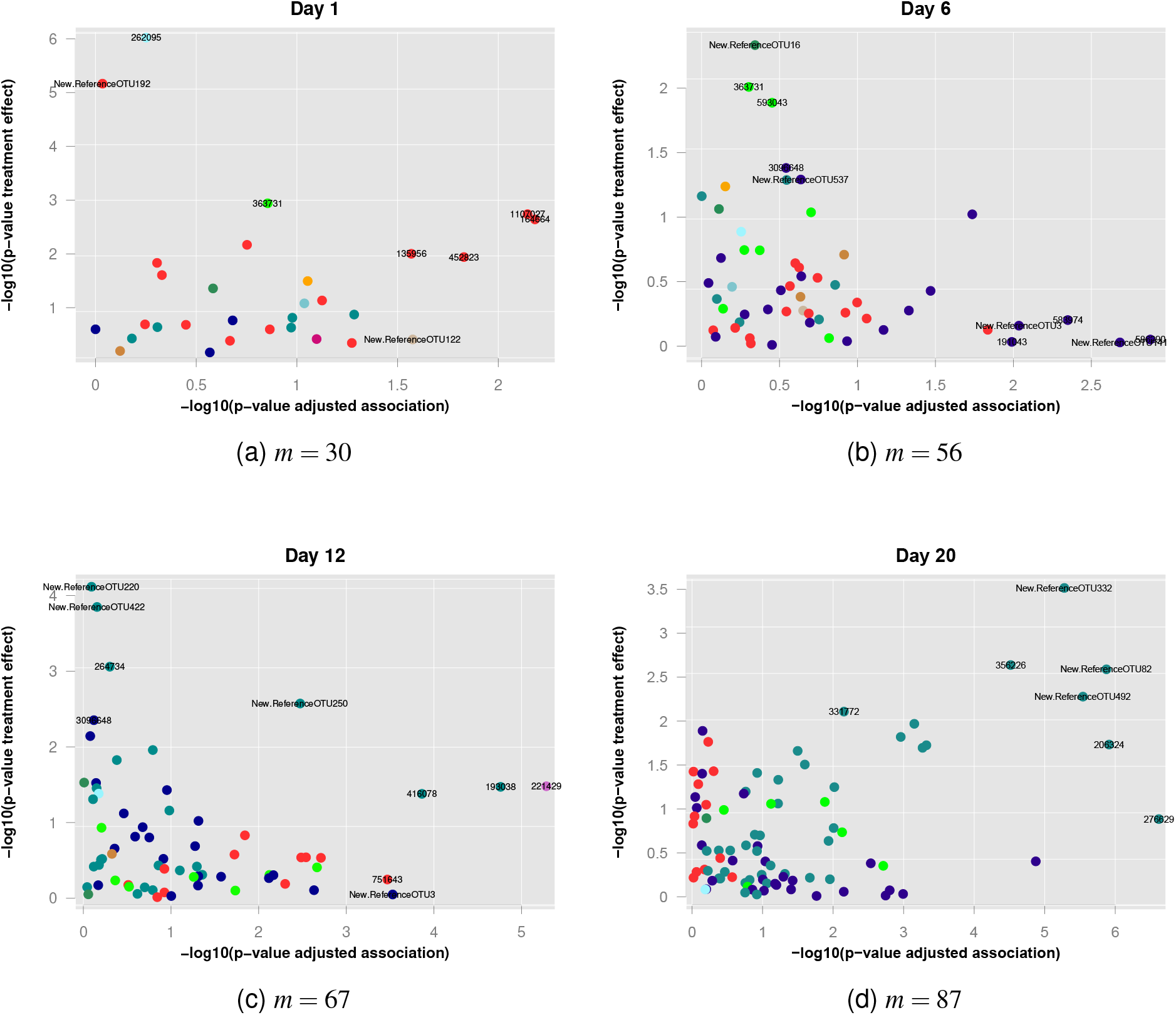
Analysis at each timepoint, −log(p-values) for adjusted association versus −log(p-values) for treatment effect.

**Figure 5:**
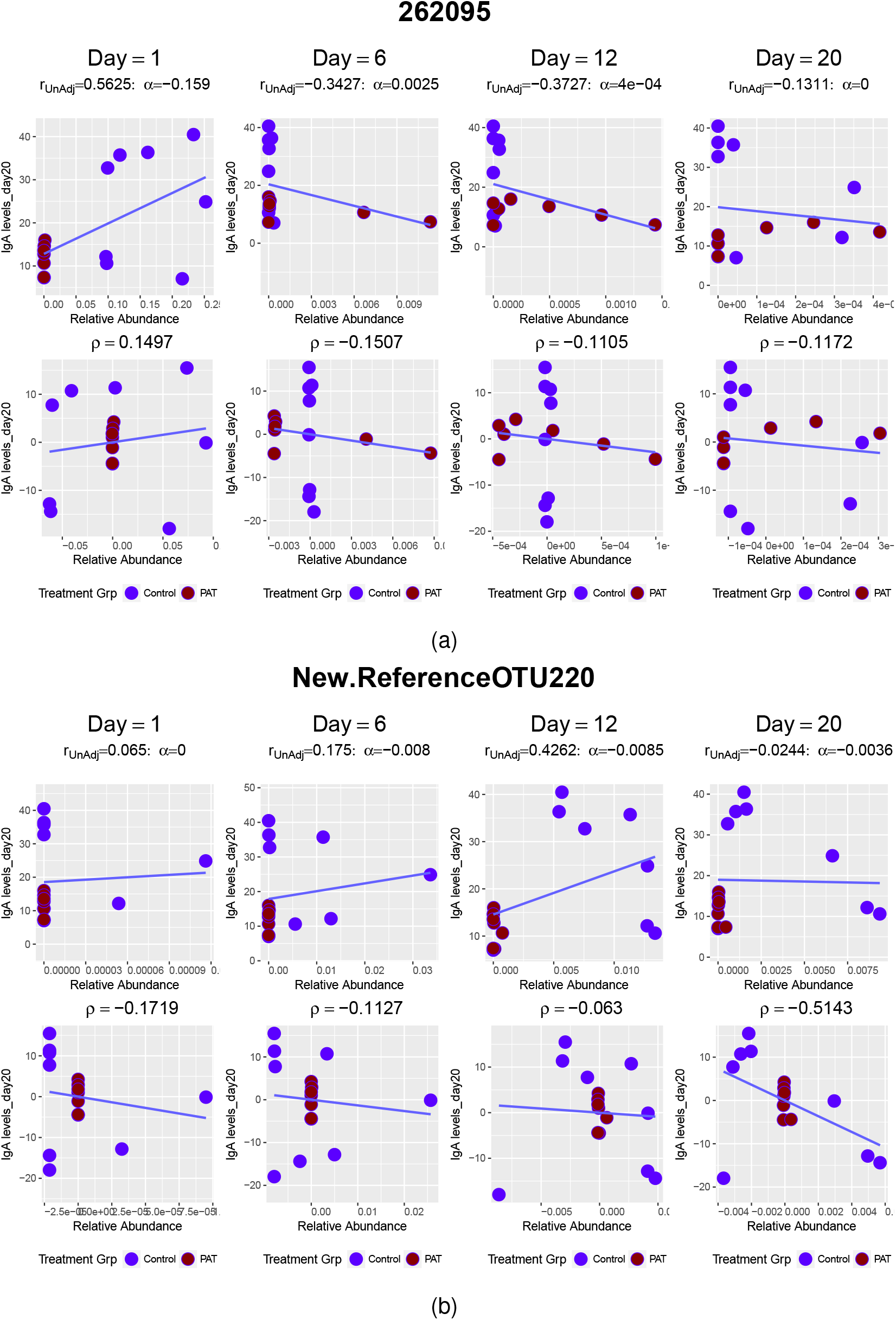
Two OTUs that are significantly (FDR = 0.05) differentially abundant - OTU 262095 and New.ReferenceOTU220 are differentially abundant at day 1 and day 12, respectively.

**Figure 6:**
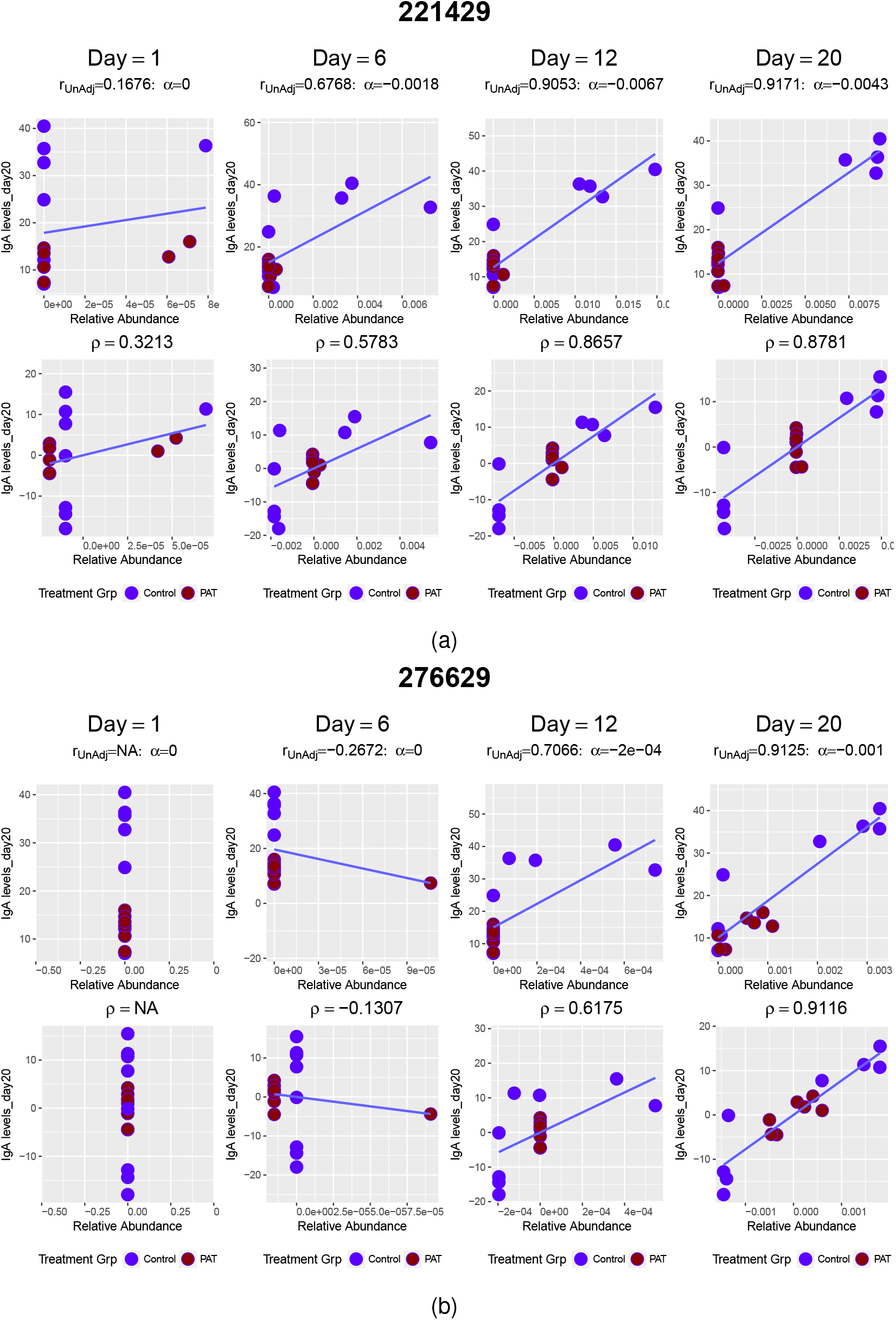
Two OTUs that are significantly (FDR = 0.05) correlated with IgA level at day 20 - OTU 221429 and OTU 276629 are significantly correlated at day 12 and day 20, respectively.

The number of differentially abundant OTUs differs from one timepoint to another. For day 1, 8 OTUs have differential sequence (with respect to relative abundance) between the PAT and Control Group (Table S3 in supplementary appendix) and most of them belongs to the *Lactobacillaceae* family. At day 6, there is no significantly differentially abundant OTU. From day 12 onwards, all the differentially abundant OTUs are from S24-7 family (cyan points in Figure 4). Moreover, as seen before (Figure 1d), OTUs from this family have higher relative abundance in the control group (negative treatment effect, Table 3) while most of the OTUs that are not from the *S24-7* family (e.g. *Bifidobacteriaceae* and *Lachnospiraceae*, marked in orange and darkblue, respectively, in FIgure 4) have higher relative abundance in the PAT group. This implies that OTUs from S24-7 family, might not be thriving in the host system with PAT-altered microbiota.

**Table 3:**
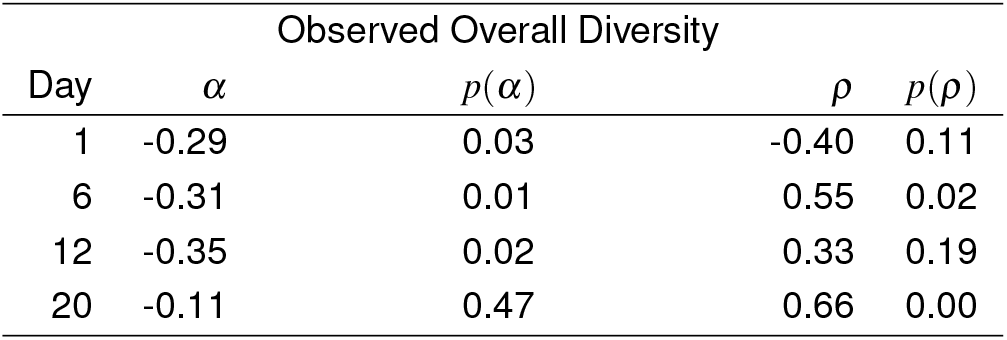
Joint Model Results for overall observed *α*-diversity and IgA (FDR = 0.05).

Figure 5 shows the 3-way association across timepoints for two of the differential OTUs. The upper row panels are the observed data and the lower panels are the data after adjusting for the treatment effect. In the scatterplot of the raw data for OTU 1107027 (Figure 5a), there is a clear separation between the two treatment groups at day 1 and hence, the treatment effect is significant for this OTU at day 1. The scenario is similar for New.ReferenceOTU220 (Figure 5b) at day 12. However, after adjusting for the treatment effect, these OTUs are not predictive of the IgA level.

Table S4 provides the estimates of the top 10 OTUs with high adjusted association by day. The OTUs from the *S24-7* family dominated at day 20. For the earlier timepoints, no OTUs were found to be significantly associated at day 1 or at day 6 while few OTUs belonging to different families can serve as predictive biomarkers at day 12 for the IgA level.

Note that no OTU is associated with the IgA level at day 1 or at day 6. However, 15 OTUs are associated at day 12 and only two of them remained associated with the IgA level until day 20. 22 other OTUs are also found to be significantly associated with IgA at day 20. New.ReferenceOTU513 is among the top 10 significantly associated OTUs both at day 12 and day 20. This OTU belongs to family *Lachnospiraceae* and is highly assoiated with the IgA level from day 12 onwards (lower panels, Figure 7) but not differentially abundant between the two groups (upper panels, Figure 7).

**Figure 7:**
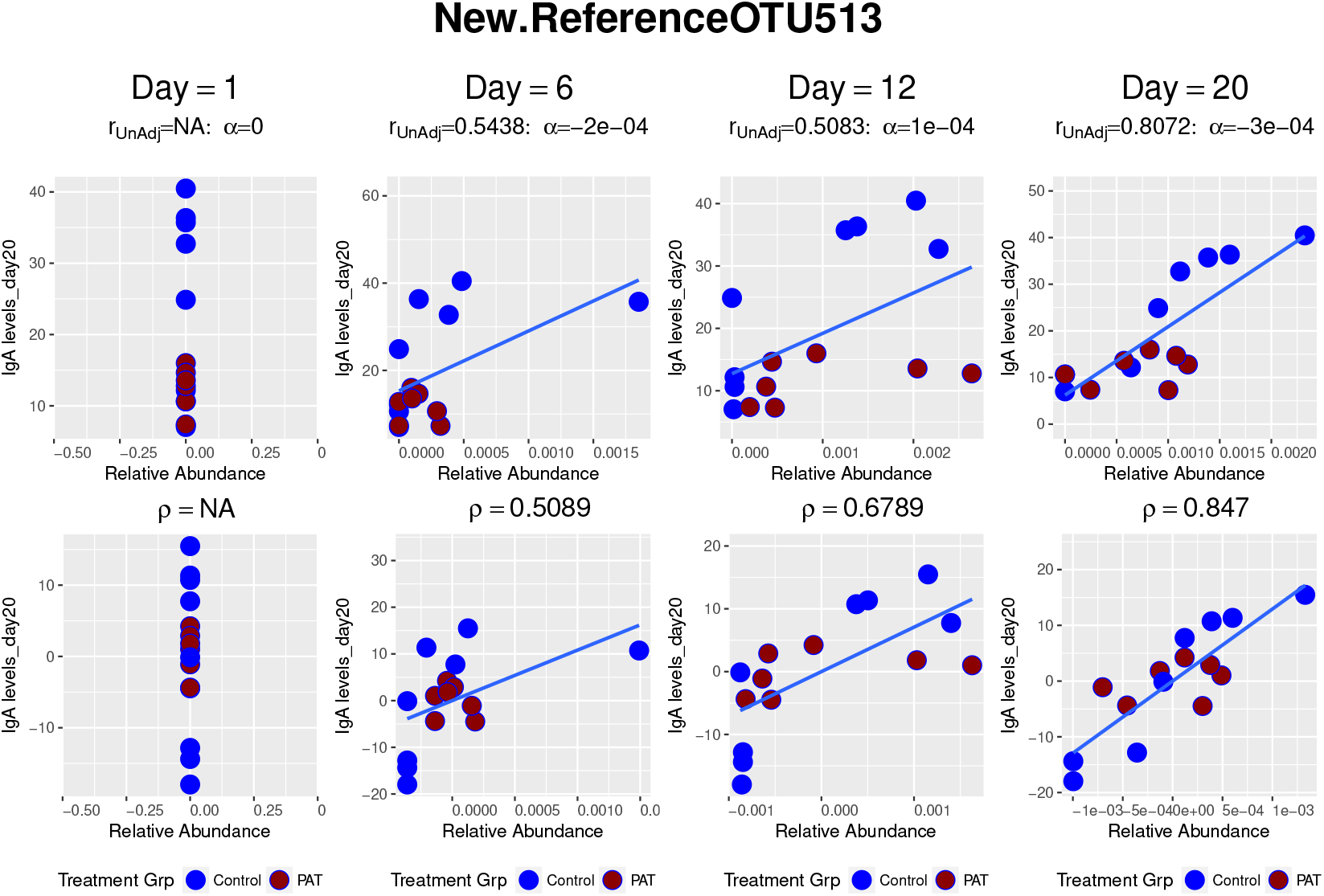
Relative abundance of NewReferenceOTU513 against IgA level over time. This OTU is found to be significantly correlated from day 12 onwards.

Based on the results of the OTU level analysis, identified OTUs from the *S24-7* are good biomarkers for the classification of treatment groups while some other OTUs (e.g OTUs from the *Lachnospiraceae* family) can serve as predictive biomarkers for the IgA level.

### 4.3 Analysis at Kingdom Level

After checking the family-level diversity, we further looked into the overall observed *α*-diversity. In this scenario, for each timepoint, our X-matrix in Figure 2 gets reduced to a vector and the other 2 variables remain similar as before.

Figure 8 and Table 3 show that in contrast with day 20, overall observed diversity is significantly differential between the two treatment groups at day 1, day 6 and day 12. When the correlation is investigated, the observed *α*-diversity is found to be significantly correlated with the IgA level at day 6 and day 20, but not at day 1 and day 12.

**Figure 8:**
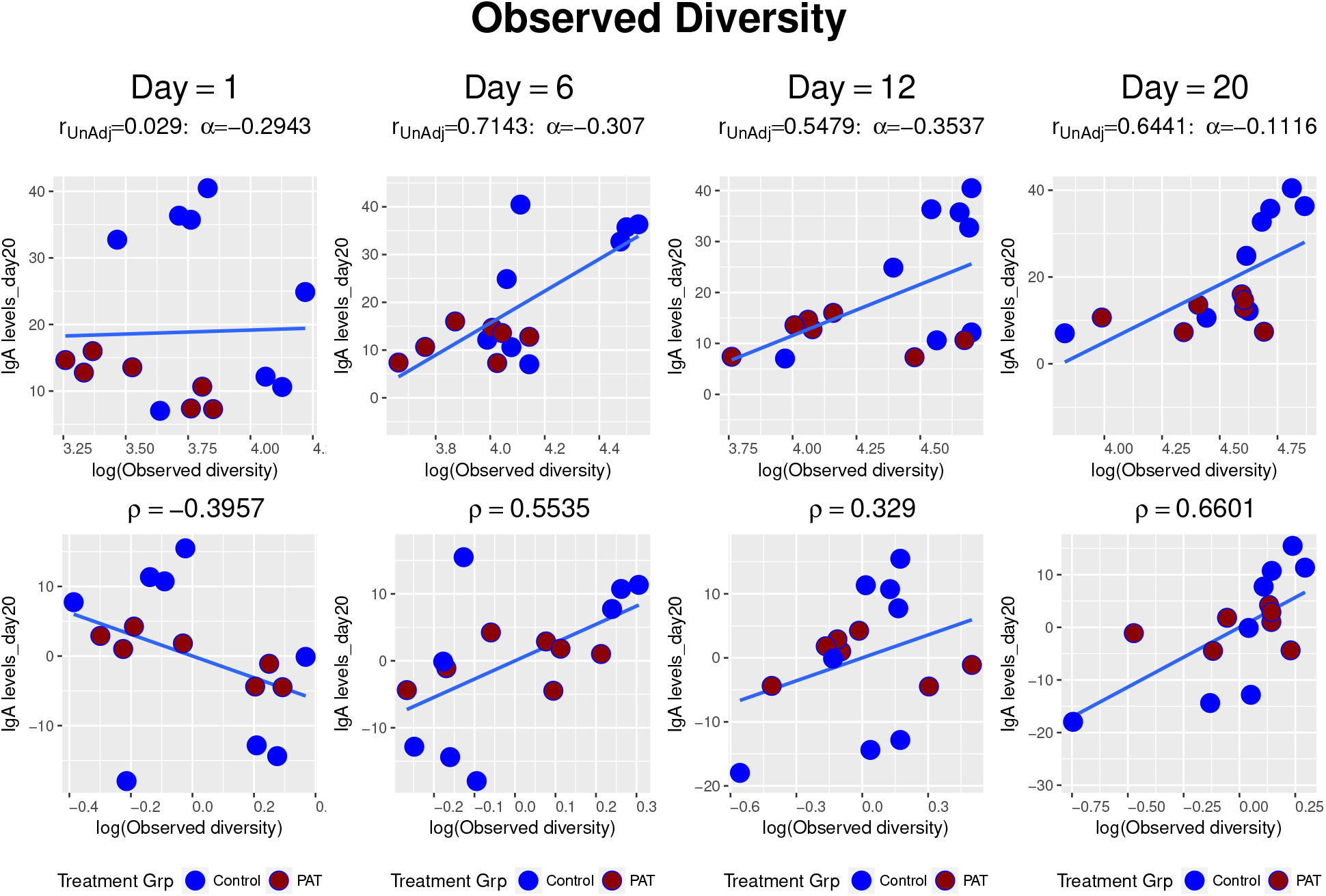
Observed diversity against IgA over time.

## 5 Discussion

In this article, the biomarker setting was introduced in the context of analysing microbiome data. The joint modeling technique can be used to identify OTUs (or families) which are differentially abundant and significantly associated with a response variable and hence, can be used as predictive biomarkers. The joint model was used at three different levels of the phylogenetic tree - OTU-level, *Family* −level and *Kingdom*-level and a certain level of thresholding is done to filter out OTUs (or families). For the OTU-level analysis relative abundance of the OTUs are used whereas different richness measures are used to do the analysis at other levels. For the discussed case study, microbiome data from earlier timepoints can also be used to predict the response at a given timepoint. After analysing the data, it was also visible that though OTUs from different families dominated at different timepoints, the *S24-7* and *Lachnospiraceae* family were found to be the most interesting ones. However, the results for the family-level richness and overall *α*-diversity differ depending on which measure of richness is being used for the analysis.

