## Supplemental Appendix File for "Development of Microbiome Biomarkers for IgA: a Joint Modeling Approach"

### 0.1 Data Processing

One of the most common characteristics of microbiome datasets is high proportion of zeroes. We perform an OTU-specific analysis and the OTUs with zero counts in at least 70% of the samples per treatment group (i.e., at least $n_{Control} = 6$ and $n_{PAT} = 5$ with zero counts) were excluded from the analysis as they were not suitable for this modeling framework. This thresholding leaves us with a total of 30, 56, 67 and 87 OTUs for day 1,6, 12, and 20 respectively (Figure S1 in Appendix). The number of active OTUs or the observed richness or $\alpha$-diversity of different samples increased over time. The highest significant difference is observed at day 12 in this case and this also motivates the investigate if microbiome from earlier timepoints affect the IgA leve at a later timepoint.

### 0.2 Results

The richness of a family is the $\alpha$-diversity, defined at family level and can be used as a measure for the microbiome activity for a family. For an experiment with $p$ families the richness matrix is given by,

$$\mathbf{F} = \left.\begin{pmatrix} f_{11} & f_{12} & \cdots & f_{1n} \\ f_{21} & f_{22} & \cdots & f_{2n} \\ . & . & . & . \\ f_{j1} & f_{j2} & \cdots & f_{jn} \\ . & . & . & . \\ f_{p1} & f_{p2} & \cdots & f_{pn} \end{pmatrix}\right\} p \text{ families} , \tag{1}$$

*CONTACT Rudradev Sengupta.

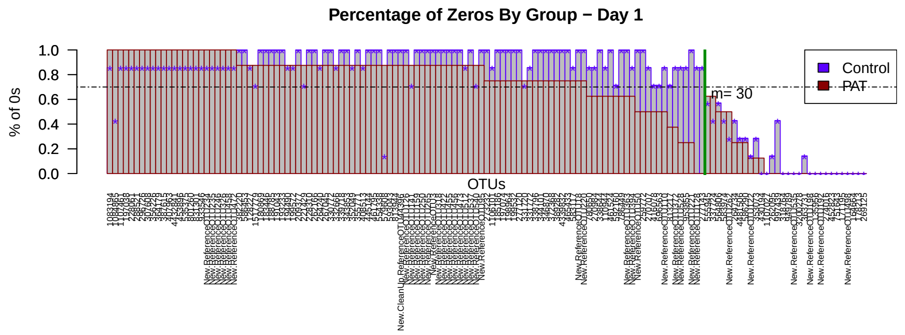


(a)

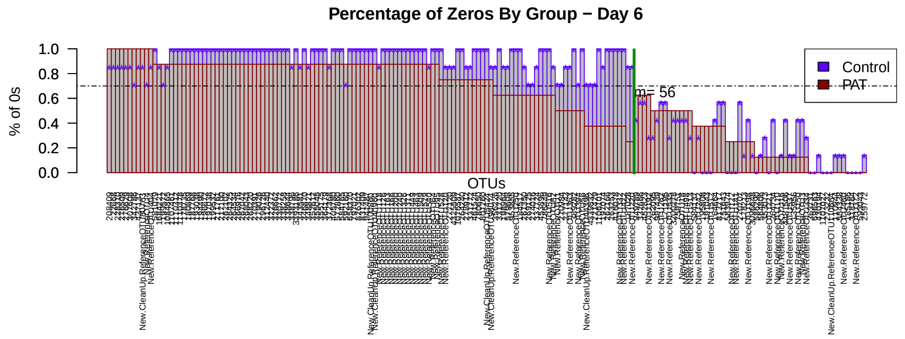


(b)

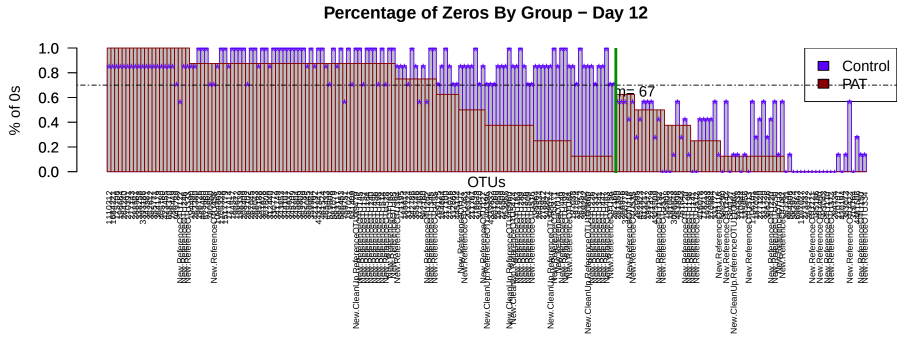


(c)

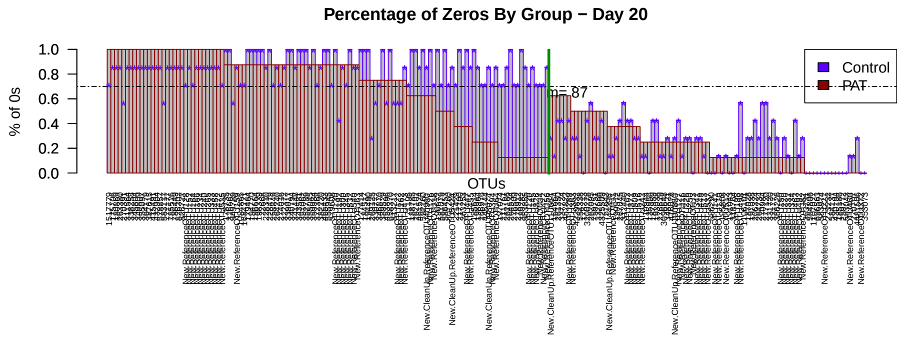


(d)

Figure S1: Barplot showing the proportion of zero counts present in each group using the 70% threshold (horizontal line). All OTU's to the left of the green line were not included in the joint model analysis.

where $f_{ji}$ is the total number of active OTUs, belonging to the $j$th family, for the $i$th sample and is calculated by

$$f_{ji} = \sum_{j' \in j} x''_{j'i}. \tag{2}$$

Here, $x''_{ji}$ is an indicator variable defined by,

$$x''_{ji} = \begin{cases} 1, & \text{if the } x_{ji} > 0, \\ 0, & \text{otherwise,} \end{cases} \tag{3}$$

where $x_{ji}$ is the count of the $j$th OTU for the $i$ subject. Figure S4 displays the observed family level richness for all the families, analyzed in the transPAT study. As mentioned earlier in the paper, Figure S4 shows that different families dominate at different timepoints. However, some families e.g., *Dehalobacteriaceae*, *Moraxellaceae* etc., are not active at any timepoint. Different measures of family level richness, Chao1 (Chao, 1984), Shannon Index (Shannon, 1948) etc. are also analyzed in a similar manner using the joint model and the results are summarized in Table S1 and Table S2, respectively. In addition, the results, corresponding to Chao1 measure and Shannon Index for overall $\alpha$-diversity, are displayed in Table S5 and Table S6, respectively.

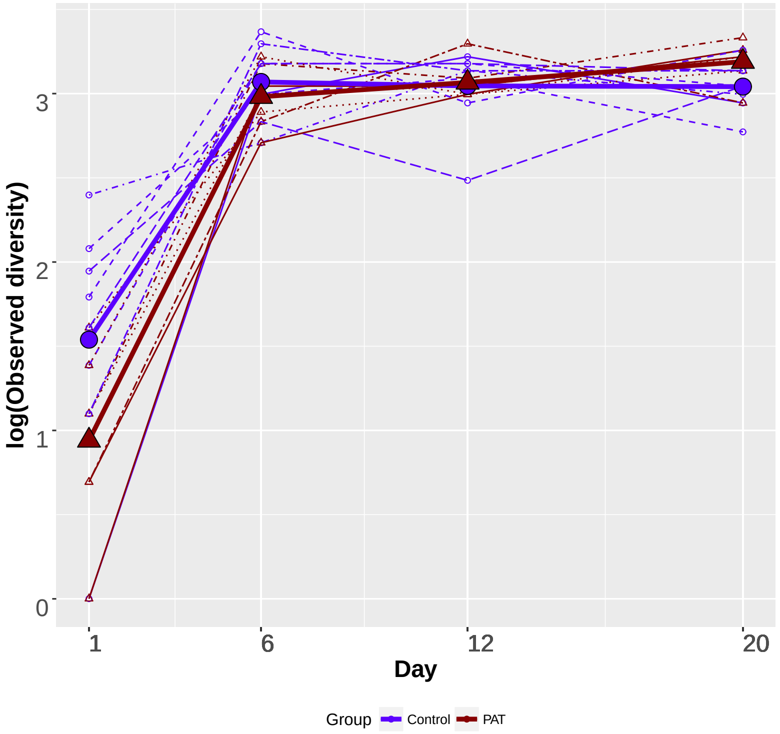


Figure S2: Individual and mean family-level diversity profiles over time for the *Lachnospiraceae* family with dashed and solid lines representing subject profiles and mean profiles, respectively.

| | | | Day 1 | | | |
|---|---|---|---|---|---|---|
| Families | $\alpha$ | $p(\alpha)$ | adj-p($\alpha$) | $\rho$ | $p(\rho)$ | adj-p($\rho$) |
| Verrucomicrobiaceae | -0.97 | 0.00 | 0.00 | -0.32 | 0.21 | 0.67 |
| Erysipelotrichaceae | -0.99 | 0.00 | 0.00 | -0.22 | 0.38 | 0.67 |
| Bifidobacteriaceae | -0.00 | 0.03 | 0.10 | -0.04 | 0.89 | 0.89 |
| Lachnospiraceae | -0.82 | 0.08 | 0.21 | 0.07 | 0.79 | 0.87 |
| Lactobacillaceae | -0.11 | 0.13 | 0.26 | -0.26 | 0.31 | 0.67 |
| S24-7 | -0.62 | 0.19 | 0.32 | -0.28 | 0.28 | 0.67 |
| Enterobacteriaceae | -0.25 | 0.28 | 0.40 | 0.43 | 0.08 | 0.67 |
| Unknown | -0.19 | 0.59 | 0.74 | -0.18 | 0.49 | 0.70 |
| Ruminococcaceae | -0.10 | 0.74 | 0.82 | -0.15 | 0.56 | 0.70 |
| Turicibacteraceae | -0.00 | 0.93 | 0.93 | -0.22 | 0.40 | 0.67 |
| | | | Day 6 | | | |
| Families | $\alpha$ | $p(\alpha)$ | adj-p($\alpha$) | $\rho$ | $p(\rho)$ | adj-p($\rho$) |
| S24-7 | -0.95 | 0.01 | 0.15 | 0.10 | 0.70 | 0.78 |
| Turicibacteraceae | 0.00 | 0.17 | 0.60 | 0.61 | 0.01 | 0.04 |
| Ruminococcaceae | -0.35 | 0.23 | 0.60 | 0.16 | 0.54 | 0.77 |
| Lachnospiraceae | -0.18 | 0.27 | 0.60 | 0.46 | 0.06 | 0.19 |
| Unknown | -0.27 | 0.34 | 0.60 | 0.27 | 0.30 | 0.59 |
| Lactobacillaceae | 0.23 | 0.42 | 0.60 | 0.62 | 0.01 | 0.04 |
| Enterococcaceae | 0.00 | 0.48 | 0.60 | 0.22 | 0.38 | 0.64 |
| Verrucomicrobiaceae | 0.03 | 0.48 | 0.60 | 0.00 | 0.99 | 0.99 |
| Erysipelotrichaceae | -0.12 | 0.63 | 0.70 | -0.10 | 0.70 | 0.78 |
| Bifidobacteriaceae | 0.00 | 0.80 | 0.80 | 0.38 | 0.13 | 0.31 |
| | | | Day 12 | | | |
| Families | $\alpha$ | $p(\alpha)$ | adj-p($\alpha$) | $\rho$ | $p(\rho)$ | adj-p($\rho$) |
| S24-7 | -0.50 | 0.08 | 0.36 | 0.40 | 0.10 | 0.27 |
| Erysipelotrichaceae | 0.26 | 0.12 | 0.36 | 0.01 | 0.97 | 0.97 |
| Verrucomicrobiaceae | 0.03 | 0.48 | 0.62 | -0.16 | 0.54 | 0.65 |
| Unknown | 0.18 | 0.53 | 0.62 | 0.56 | 0.02 | 0.10 |
| Lachnospiraceae | -0.08 | 0.56 | 0.62 | -0.34 | 0.18 | 0.27 |
| Lactobacillaceae | 0.09 | 0.62 | 0.62 | 0.34 | 0.17 | 0.27 |
| | | | Day 20 | | | |
| Families | $\alpha$ | $p(\alpha)$ | adj-p($\alpha$) | $\rho$ | $p(\rho)$ | adj-p($\rho$) |
| Lachnospiraceae | 0.21 | 0.01 | 0.07 | 0.56 | 0.02 | 0.06 |
| Erysipelotrichaceae | 0.20 | 0.11 | 0.43 | 0.11 | 0.67 | 0.68 |
| Lactobacillaceae | 0.20 | 0.32 | 0.85 | 0.31 | 0.21 | 0.34 |
| Ruminococcaceae | -0.13 | 0.69 | 0.92 | 0.54 | 0.02 | 0.06 |
| Verrucomicrobiaceae | -0.01 | 0.73 | 0.92 | -0.11 | 0.68 | 0.68 |
| S24-7 | -0.09 | 0.84 | 0.92 | 0.52 | 0.03 | 0.06 |
| Unknown | -0.07 | 0.84 | 0.92 | 0.61 | 0.01 | 0.06 |
| Rikenellaceae | -0.03 | 0.92 | 0.92 | 0.20 | 0.44 | 0.58 |

Table S1: Parameter estimates from the model for all the families at different timepoints with respect to log(Chao1) measure for observed diversity. Results are sorted according to the adjusted p-values for the treatment effect $\alpha$.

| Day 1 | | | | | | |
|---|---|---|---|---|---|---|
| Families | $\alpha$ | $p(\alpha)$ | adj-p($\alpha$) | $\rho$ | $p(\rho)$ | adj-p($\rho$) |
| Lactobacillaceae | -0.04 | 0.56 | 0.67 | -0.59 | 0.01 | 0.03 |
| Lachnospiraceae | -0.14 | 0.60 | 0.67 | 0.44 | 0.07 | 0.10 |
| S24-7 | -0.14 | 0.67 | 0.67 | -0.20 | 0.44 | 0.44 |

| Day 6 | | | | | | |
|---|---|---|---|---|---|---|
| Families | $\alpha$ | $p(\alpha)$ | adj-p($\alpha$) | $\rho$ | $p(\rho)$ | adj-p($\rho$) |
| Unknown | -0.19 | 0.16 | 0.56 | -0.16 | 0.52 | 0.53 |
| S24-7 | -0.35 | 0.19 | 0.56 | 0.42 | 0.09 | 0.53 |
| Lactobacillaceae | 0.05 | 0.29 | 0.59 | 0.32 | 0.20 | 0.53 |
| Verrucomicrobiaceae | 0.01 | 0.71 | 0.94 | -0.18 | 0.49 | 0.53 |
| Enterobacteriaceae | -0.04 | 0.78 | 0.94 | -0.21 | 0.42 | 0.53 |
| Lachnospiraceae | 0.01 | 0.96 | 0.96 | 0.16 | 0.53 | 0.53 |

| Day 12 | | | | | | |
|---|---|---|---|---|---|---|
| Families | $\alpha$ | $p(\alpha)$ | adj-p($\alpha$) | $\rho$ | $p(\rho)$ | adj-p($\rho$) |
| S24-7 | -1.11 | 0.00 | 0.00 | -0.44 | 0.07 | 0.18 |
| Lactobacillaceae | 0.04 | 0.20 | 0.50 | 0.09 | 0.74 | 0.88 |
| Unknown | -0.16 | 0.30 | 0.51 | 0.57 | 0.01 | 0.07 |
| Verrucomicrobiaceae | 0.00 | 0.53 | 0.66 | 0.37 | 0.14 | 0.23 |
| Lachnospiraceae | -0.01 | 0.94 | 0.94 | -0.04 | 0.88 | 0.88 |

| Day 20 | | | | | | |
|---|---|---|---|---|---|---|
| Families | $\alpha$ | $p(\alpha)$ | adj-p($\alpha$) | $\rho$ | $p(\rho)$ | adj-p($\rho$) |
| S24-7 | -0.64 | 0.01 | 0.02 | 0.29 | 0.26 | 0.54 |
| Lachnospiraceae | 0.27 | 0.04 | 0.08 | 0.02 | 0.94 | 0.94 |
| Lactobacillaceae | 0.02 | 0.69 | 0.87 | 0.21 | 0.40 | 0.54 |
| Verrucomicrobiaceae | -0.00 | 0.87 | 0.87 | 0.22 | 0.39 | 0.54 |

Table S2: Parameter estimates from the model for all the families at different timepoints with respect tothe Shannon index for observed diversity. Results are sorted according to the adjusted p-values for the treatment effect $\alpha$.

| Day 1 | | | | | | | |
|---|---|---|---|---|---|---|---|
| OTUs | $\alpha$ | $p(\alpha)$ | adj-p($\alpha$) | $\rho$ | $p(\rho)$ | adj-p($\rho$) | Family |
| 262095 | -0.16 | 0.00 | 0.00 | 0.15 | 0.56 | 0.66 | Erysipelotrichaceae |
| New.ReferenceOTU192 | 0.01 | 0.00 | 0.00 | 0.03 | 0.92 | 0.95 | Lactobacillaceae |
| 363731 | -0.01 | 0.00 | 0.01 | -0.37 | 0.14 | 0.28 | Verrucomicrobiaceae |
| 1107027 | 0.24 | 0.00 | 0.01 | 0.62 | 0.01 | 0.11 | Lactobacillaceae |
| 164664 | 0.00 | 0.00 | 0.01 | 0.62 | 0.01 | 0.11 | Lactobacillaceae |
| New.ReferenceOTU498 | 0.00 | 0.01 | 0.03 | 0.34 | 0.18 | 0.33 | Lactobacillaceae |
| 135956 | 0.00 | 0.01 | 0.04 | 0.53 | 0.03 | 0.16 | Lactobacillaceae |
| 452823 | 0.00 | 0.01 | 0.04 | 0.57 | 0.01 | 0.15 | Lactobacillaceae |
| 171195 | 0.00 | 0.01 | 0.05 | 0.17 | 0.49 | 0.62 | Lactobacillaceae |
| New.ReferenceOTU198 | 0.00 | 0.02 | 0.07 | 0.19 | 0.47 | 0.62 | Lactobacillaceae |

| Day 6 | | | | | | | |
|---|---|---|---|---|---|---|---|
| OTUs | $\alpha$ | $p(\alpha)$ | adj-p($\alpha$) | $\rho$ | $p(\rho)$ | adj-p($\rho$) | Family |
| New.ReferenceOTU16 | 0.00 | 0.00 | 0.24 | 0.19 | 0.45 | 0.67 | |
| 363731 | 0.22 | 0.01 | 0.24 | 0.17 | 0.50 | 0.68 | Verrucomicrobiaceae |
| 593043 | 0.00 | 0.01 | 0.24 | 0.24 | 0.35 | 0.57 | Verrucomicrobiaceae |
| 3096648 | 0.00 | 0.04 | 0.46 | 0.27 | 0.29 | 0.50 | Lachnospiraceae |
| New.ReferenceOTU537 | 0.00 | 0.05 | 0.46 | 0.30 | 0.23 | 0.49 | Lachnospiraceae |
| 430194 | -0.15 | 0.05 | 0.46 | -0.27 | 0.28 | 0.50 | S24-7 |
| 997439 | 0.00 | 0.06 | 0.46 | 0.10 | 0.70 | 0.82 | Bifidobacteriaceae |
| 264734 | -0.06 | 0.07 | 0.48 | -0.00 | 1.00 | 1.00 | S24-7 |
| New.ReferenceOTU535 | 0.00 | 0.09 | 0.48 | 0.07 | 0.77 | 0.85 | |
| 185186 | 0.00 | 0.09 | 0.48 | 0.32 | 0.20 | 0.49 | Verrucomicrobiaceae |

| Day 12 | | | | | | | |
|---|---|---|---|---|---|---|---|
| OTUs | $\alpha$ | $p(\alpha)$ | adj-p($\alpha$) | $\rho$ | $p(\rho)$ | adj-p($\rho$) | Family |
| New.ReferenceOTU220 | -0.01 | 0.00 | 0.00 | -0.06 | 0.81 | 0.86 | S24-7 |
| New.ReferenceOTU422 | -0.00 | 0.00 | 0.00 | -0.10 | 0.70 | 0.81 | S24-7 |
| 264734 | -0.06 | 0.00 | 0.02 | -0.17 | 0.50 | 0.67 | S24-7 |
| New.ReferenceOTU250 | -0.01 | 0.00 | 0.04 | 0.66 | 0.00 | 0.02 | S24-7 |
| 3096648 | 0.00 | 0.00 | 0.06 | 0.08 | 0.76 | 0.84 | Lachnospiraceae |
| New.ReferenceOTU537 | 0.00 | 0.01 | 0.08 | 0.05 | 0.83 | 0.87 | Lachnospiraceae |
| 1106101 | -0.01 | 0.01 | 0.11 | 0.35 | 0.16 | 0.29 | S24-7 |
| New.ReferenceOTU82 | -0.00 | 0.01 | 0.13 | 0.21 | 0.42 | 0.60 | S24-7 |
| New.ReferenceOTU16 | 0.00 | 0.03 | 0.17 | 0.01 | 0.98 | 0.98 | |
| 259772 | 0.00 | 0.03 | 0.17 | 0.09 | 0.72 | 0.81 | Lachnospiraceae |

| Day 20 | | | | | | | |
|---|---|---|---|---|---|---|---|
| OTUs | $\alpha$ | $p(\alpha)$ | adj-p($\alpha$) | $\rho$ | $p(\rho)$ | adj-p($\rho$) | Family |
| New.ReferenceOTU332 | -0.00 | 0.00 | 0.03 | 0.87 | 0.00 | 0.00 | S24-7 |
| 356226 | -0.01 | 0.00 | 0.07 | 0.83 | 0.00 | 0.00 | S24-7 |
| New.ReferenceOTU82 | -0.01 | 0.00 | 0.07 | 0.89 | 0.00 | 0.00 | S24-7 |
| New.ReferenceOTU492 | -0.00 | 0.01 | 0.11 | 0.88 | 0.00 | 0.00 | S24-7 |
| 331772 | -0.01 | 0.01 | 0.14 | 0.62 | 0.01 | 0.03 | S24-7 |
| 353073 | -0.13 | 0.01 | 0.15 | 0.73 | 0.00 | 0.01 | S24-7 |
| 259772 | 0.00 | 0.01 | 0.15 | -0.10 | 0.71 | 0.80 | Lachnospiraceae |
| 331720 | -0.01 | 0.02 | 0.15 | 0.71 | 0.00 | 0.01 | S24-7 |
| 178213 | 0.00 | 0.02 | 0.15 | -0.14 | 0.59 | 0.74 | Lactobacillaceae |
| 206324 | -0.00 | 0.02 | 0.15 | 0.89 | 0.00 | 0.00 | S24-7 |

Table S3: Parameter estimates from the model for top 10 differentially abundant OTUs (FDR = 0.05) at different timepoints. The results are sorted according to the adjusted p-values for the treatment effect $\alpha$.

| Day 1 | | | | | | | |
|---|---|---|---|---|---|---|---|
| OTUs | $\alpha$ | $p(\alpha)$ | adj-p($\alpha$) | $\rho$ | $p(\rho)$ | adj-p($\rho$) | Family |
| 1107027 | 0.24 | 0.00 | 0.01 | 0.62 | 0.01 | 0.11 | Lactobacillaceae |
| 164664 | 0.00 | 0.00 | 0.01 | 0.62 | 0.01 | 0.11 | Lactobacillaceae |
| 452823 | 0.00 | 0.01 | 0.04 | 0.57 | 0.01 | 0.15 | Lactobacillaceae |
| 135956 | 0.00 | 0.01 | 0.04 | 0.53 | 0.03 | 0.16 | Lactobacillaceae |
| New.ReferenceOTU122 | -0.03 | 0.39 | 0.45 | -0.53 | 0.03 | 0.16 | Turicibacteraceae |
| 353073 | -0.00 | 0.13 | 0.27 | -0.47 | 0.05 | 0.23 | S24-7 |
| 274016 | -0.00 | 0.45 | 0.49 | -0.47 | 0.05 | 0.23 | Lactobacillaceae |
| 997439 | -0.03 | 0.03 | 0.09 | -0.42 | 0.09 | 0.25 | Bifidobacteriaceae |
| 178213 | 0.00 | 0.07 | 0.17 | 0.44 | 0.07 | 0.25 | Lactobacillaceae |
| 949789 | 0.01 | 0.08 | 0.18 | -0.42 | 0.09 | 0.25 | Enterococcaceae |

| Day 6 | | | | | | | |
|---|---|---|---|---|---|---|---|
| OTUs | $\alpha$ | $p(\alpha)$ | adj-p($\alpha$) | $\rho$ | $p(\rho)$ | adj-p($\rho$) | Family |
| 586290 | -0.00 | 0.89 | 0.97 | 0.71 | 0.00 | 0.06 | Lachnospiraceae |
| New.ReferenceOTU141 | -0.00 | 0.94 | 0.97 | 0.68 | 0.00 | 0.06 | Lachnospiraceae |
| 583974 | -0.00 | 0.63 | 0.88 | 0.65 | 0.00 | 0.08 | Lachnospiraceae |
| New.ReferenceOTU3 | 0.00 | 0.69 | 0.90 | 0.60 | 0.01 | 0.11 | Lachnospiraceae |
| 191043 | -0.00 | 0.93 | 0.97 | 0.60 | 0.01 | 0.11 | Lachnospiraceae |
| New.ReferenceOTU512 | 0.00 | 0.75 | 0.90 | 0.57 | 0.01 | 0.14 | Lactobacillaceae |
| 584137 | 0.00 | 0.10 | 0.48 | 0.56 | 0.02 | 0.15 | Lachnospiraceae |
| New.ReferenceOTU513 | -0.00 | 0.37 | 0.80 | 0.51 | 0.03 | 0.24 | Lachnospiraceae |
| 564806 | -0.02 | 0.53 | 0.86 | -0.48 | 0.05 | 0.29 | Lachnospiraceae |
| 195808 | -0.00 | 0.75 | 0.90 | 0.45 | 0.07 | 0.38 | Lachnospiraceae |

| Day 12 | | | | | | | |
|---|---|---|---|---|---|---|---|
| OTUs | $\alpha$ | $p(\alpha)$ | adj-p($\alpha$) | $\rho$ | $p(\rho)$ | adj-p($\rho$) | Family |
| 221429 | -0.01 | 0.03 | 0.17 | 0.87 | 0.00 | 0.00 | Ruminococcaceae |
| 193038 | -0.00 | 0.03 | 0.17 | 0.84 | 0.00 | 0.00 | S24-7 |
| 416078 | -0.00 | 0.04 | 0.17 | 0.79 | 0.00 | 0.00 | S24-7 |
| 751643 | -0.00 | 0.57 | 0.76 | 0.76 | 0.00 | 0.00 | Lactobacillaceae |
| New.ReferenceOTU3 | -0.00 | 0.89 | 0.92 | 0.76 | 0.00 | 0.00 | Lachnospiraceae |
| 318764 | -0.00 | 0.29 | 0.59 | 0.69 | 0.00 | 0.02 | Lactobacillaceae |
| 185186 | 0.00 | 0.39 | 0.66 | -0.68 | 0.00 | 0.02 | Verrucomicrobiaceae |
| New.ReferenceOTU513 | 0.00 | 0.78 | 0.87 | 0.68 | 0.00 | 0.02 | Lachnospiraceae |
| New.ReferenceOTU250 | -0.01 | 0.00 | 0.04 | 0.66 | 0.00 | 0.02 | S24-7 |
| New.ReferenceOTU198 | 0.00 | 0.29 | 0.59 | 0.67 | 0.00 | 0.02 | Lactobacillaceae |

| Day 20 | | | | | | | |
|---|---|---|---|---|---|---|---|
| OTUs | $\alpha$ | $p(\alpha)$ | adj-p($\alpha$) | $\rho$ | $p(\rho)$ | adj-p($\rho$) | Family |
| 276629 | -0.00 | 0.13 | 0.34 | 0.91 | 0.00 | 0.00 | S24-7 |
| New.ReferenceOTU82 | -0.01 | 0.00 | 0.07 | 0.89 | 0.00 | 0.00 | S24-7 |
| 206324 | -0.00 | 0.02 | 0.15 | 0.89 | 0.00 | 0.00 | S24-7 |
| New.ReferenceOTU492 | -0.00 | 0.01 | 0.11 | 0.88 | 0.00 | 0.00 | S24-7 |
| New.ReferenceOTU332 | -0.00 | 0.00 | 0.03 | 0.87 | 0.00 | 0.00 | S24-7 |
| New.ReferenceOTU513 | -0.00 | 0.39 | 0.69 | 0.85 | 0.00 | 0.00 | Lachnospiraceae |
| 356226 | -0.01 | 0.00 | 0.07 | 0.83 | 0.00 | 0.00 | S24-7 |
| New.ReferenceOTU127 | -0.00 | 0.02 | 0.15 | 0.75 | 0.00 | 0.01 | S24-7 |
| 277120 | -0.00 | 0.02 | 0.15 | 0.74 | 0.00 | 0.01 | S24-7 |
| 353073 | -0.13 | 0.01 | 0.15 | 0.73 | 0.00 | 0.01 | S24-7 |

Table S4: Parameter estimates from the model for top 10 associated OTUs (FDR = 0.05) with IgA at different timepoints. The results are sorted according to the adjusted p-values for the adjusted association $\rho$.

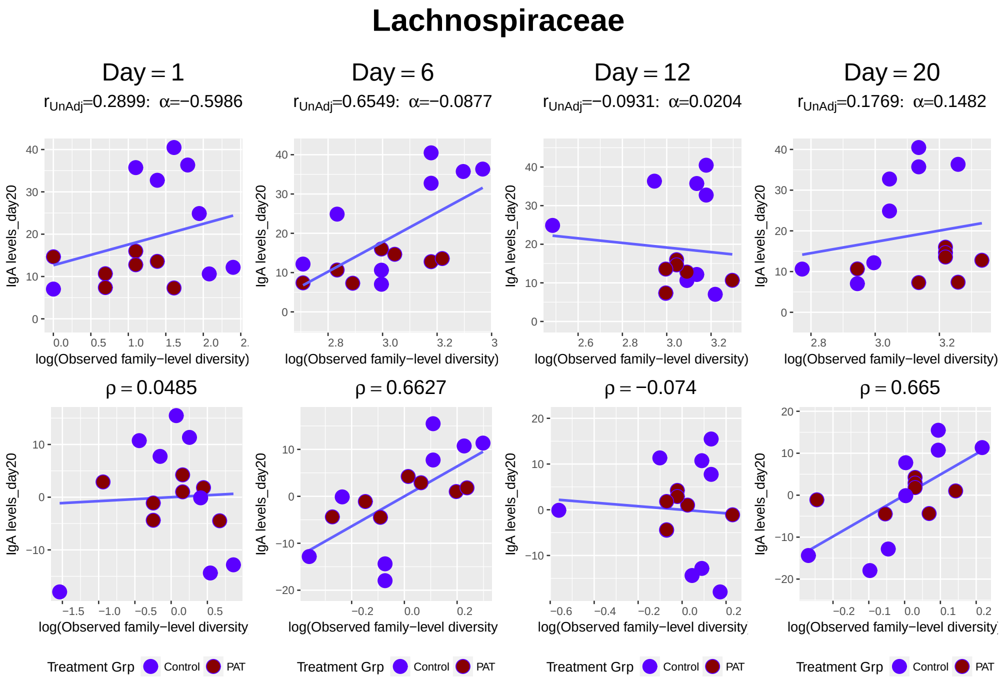


Figure S3: *Lachnospiraceae* family against IgA level over time.

| Chao1 measure for Overall Diversity | | | | |
|---|---|---|---|---|
| Day | $\alpha$ | $p(\alpha)$ | $\rho$ | $p(\rho)$ |
| 1 | -0.42 | 0.09 | -0.34 | 0.18 |
| 6 | -0.23 | 0.15 | 0.51 | 0.03 |
| 12 | -0.31 | 0.05 | -0.01 | 0.98 |
| 20 | 0.03 | 0.82 | 0.68 | 0.00 |

Table S5: Joint Model Results for log(Chao1) measure of overall observed $\alpha$-diversity and IgA (FDR = 0.05).

| Shannon Index for Overall Diversity | | | | |
|---|---|---|---|---|
| Day | $\alpha$ | $p(\alpha)$ | $\rho$ | $p(\rho)$ |
| 1 | -0.62 | 0.00 | -0.63 | 0.01 |
| 6 | -0.42 | 0.01 | 0.06 | 0.81 |
| 12 | -0.63 | 0.01 | 0.46 | 0.06 |
| 20 | -0.60 | 0.01 | 0.61 | 0.01 |

Table S6: Joint Model Results for Shannon Index of overall observed $\alpha$-diversity and IgA (FDR = 0.05).

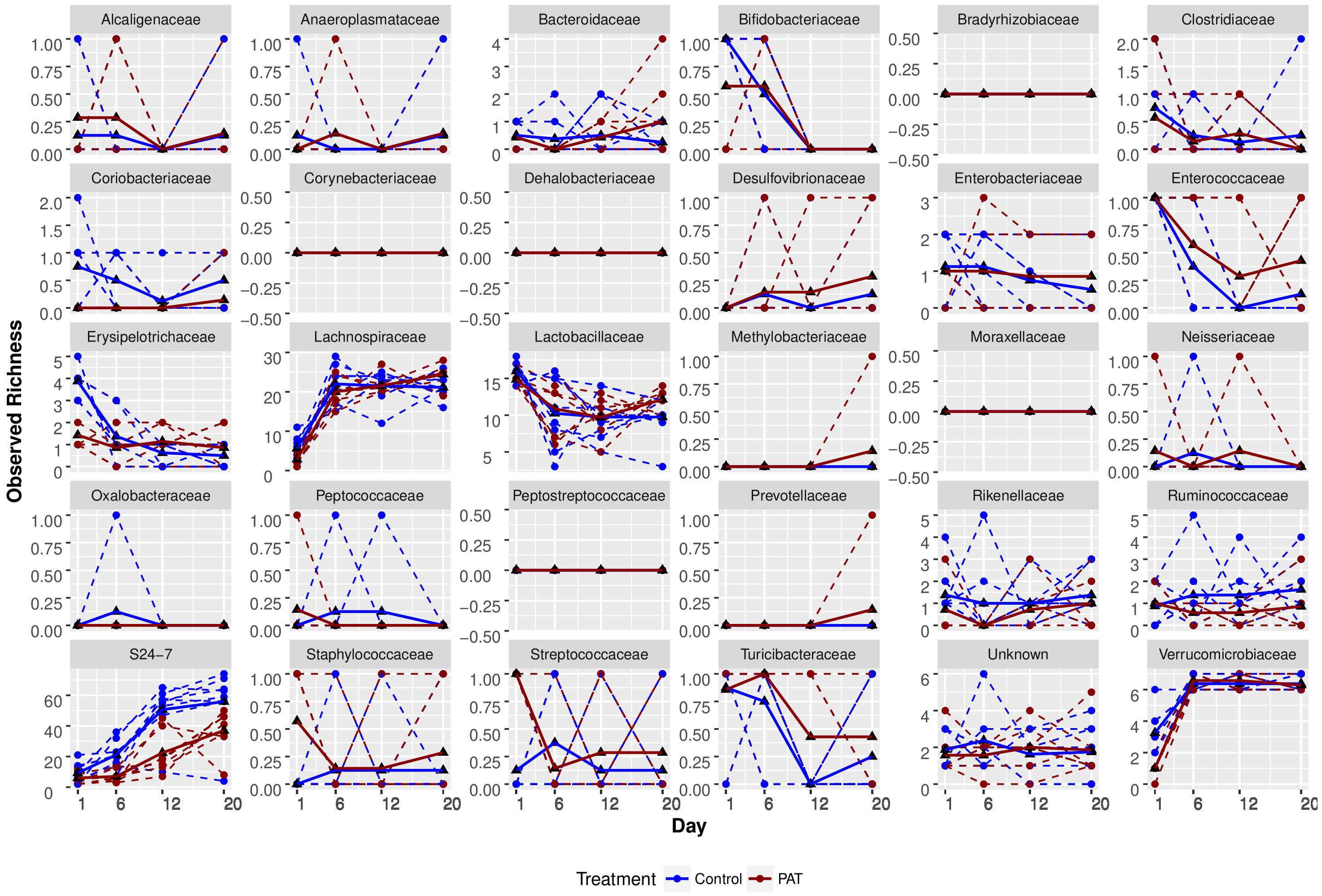


Figure S4: Observed richness for all the families over time Observed richness for all the families over time with dashed and solid lines representing subject mean profiles, respectively.

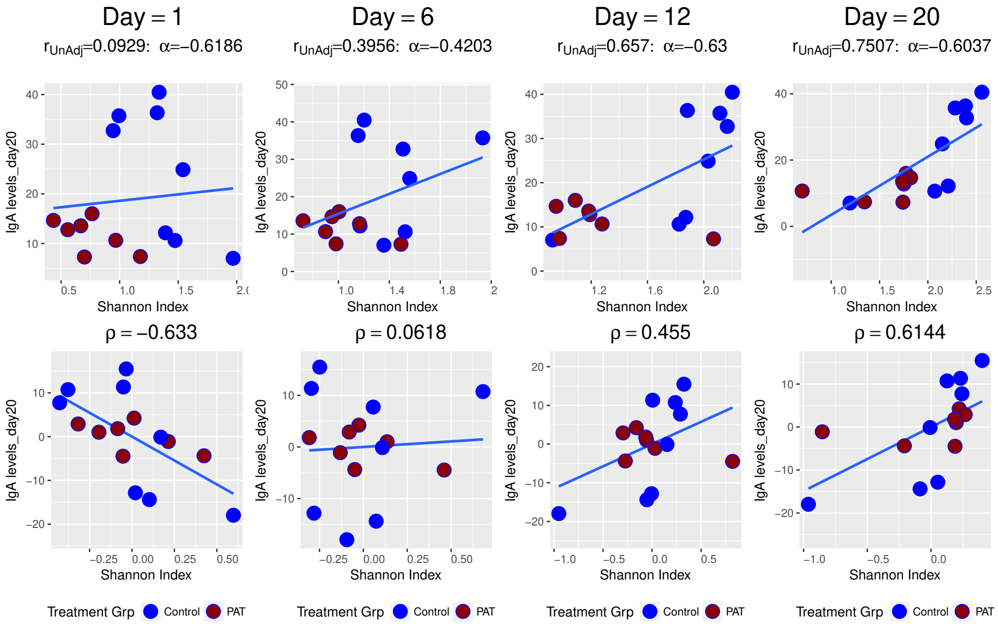


Figure S5: Shannon index for overall $\alpha$ - diversity against IgA level over time.
